# SPOT-Contact-Single: Improving Single-Sequence-Based Prediction of Protein Contact Map using a Transformer Language Model

**DOI:** 10.1101/2021.06.19.449089

**Authors:** Jaspreet Singh, Thomas Litfin, Jaswinder Singh, Kuldip Paliwal, Yaoqi Zhou

## Abstract

**Motivation:** Accurate prediction of protein contact-map is essential for accurate protein structure and function prediction. As a result, many methods have been developed for protein contact map prediction. However, most methods rely on protein-sequence-evolutionary information, which may not exist for many proteins due to lack of naturally occurring homologous sequences. Moreover, generating evolutionary profiles is computationally intensive. Here, we developed a contact-map predictor utilizing the output of a pre-trained language model ESM-1b as an input along with a large training set and an ensemble of residual neural networks.

**Results:** We showed that the proposed method makes a significant improvement over a single-sequence-based predictor SSCpred with 15% improvement in the F1-score for the independent CASP14-FM test set. It also outperforms evolutionary-profile-based methods TrRosetta and SPOT-Contact with 48.7% and 48.5% respective improvement in the F1-score on the proteins without homologs (Neff=1) in the independent SPOT-2018 set. The new method provides a much faster and reasonably accurate alternative to evolution-based methods, useful for large-scale prediction.

**Availability:** Stand-alone-version of SPOT-Contact-Single is available at https://github.com/jas-preet/SPOT-Contact-Single. Direct prediction can also be made at https://sparks-lab.org/server/spot-contact-single. The datasets used in this research can also be downloaded from the GitHub.

**Contact:**, and

**Supplementary information:** Supplementary data are available at *Bioinformatics* online.

## 1 Introduction

The past two decades have seen many developments in the field of protein structure prediction (Hanson *et al.*, 2020;Cheng *et al.*, 2019; Liu *et al.*, 2021). Significant headway has been observed specifically for protein secondary structure prediction and contact- and distance-map prediction (Hanson *et al.*, 2019; Wang *et al.*, 2016; Fang *et al.*, 2018; Wu *et al.*, 2020; Li *et al.*, 2019). These improvements have ultimately led to a considerable improvement in protein tertiary structure prediction, as observed in CASP13 (Cheng *et al.*, 2019).

Protein contact maps have been predicted by statistical inference based on Potts model and deep learning-based predictors. The predictors based on statistical inference are CCMpred (Seemayer *et al.*, 2014), Gremlin (Ovchinnikov *et al.*, 2014), EVFold (Sheridan *et al.*, 2015), plmDCA (Ekeberg *et al.*, 2014), FreeContact (Kaján *et al.*, 2014), and MetaPSICOV (Jones *et al.*, 2015). These methods were further improved by supervised deep learning-based methods such as RaptorX-Contact (Wang *et al.*, 2017), DeepCov (Jones and Kandathil, 2018), SPOT-Contact (Hanson *et al.*, 2018), and TrRosetta (Wu *et al.*, 2020).

A common trait among these methods is the use of multiple sequence alignment (MSA) and other homology-based profile information. However, many proteins have very few or no homologs to generate MSA and homology profiles (Ovchinnikov *et al.*, 2017). In this case, their performance drops significantly (Chen *et al.*, 2020). Thus, it becomes essential to develop a method that predicts protein contact maps without using homologous information.

SSCpred (Chen *et al.*, 2020) is a recently published method that predicts contact maps using the one-hot encoding of the fasta sequence and the predicted one-dimensional structural properties of SPIDER3-Single (Heffernan *et al.*, 2018). The method employs a fully convolutional model with 30 ResNet blocks. The method performs adequately for proteins with few homologs but relatively poorer for those proteins with more effective homologs when compared to MSA-based techniques (Chen *et al.*, 2020). This limitation is expected as single-sequence-based method provides less information for the neural network to learn.

To improve the performance of single-sequence-based methods for the proteins with few homologs, there is a need for exploring other possible features beyond one-hot encoding. Recently, unsupervised deep learning methods were introduced to extract features inspired by Natural Language Processing’s (NLP) language models (LM) (Rao *et al.*, 2019; Heinzinger *et al.*, 2019; Elnaggar *et al.*, 2020; Rao *et al.*, 2020). These methods are trained on protein reference libraries such as Uniref (Suzek *et al.*, 2007), Uniclust (Mirdita *et al.*, 2017), Pfam (Bateman *et al.*, 2004), and BFD (Steinegger *et al.*, 2019b; Steinegger and Söding, 2018). One unsupervised learning method is ESM-1b (Rao *et al.*, 2020). This method uses a Transformer-34 model trained on Uniref50 and outputs an embedding and attention maps as output (Rao *et al.*, 2020).

In this work, we examined the use of ESM-1b’s attention map as an input feature to our model to improve the contact-map prediction of our single-sequence-based method. We demonstrated that unsupervised learning features concatenated with one-hot encoding and SPOT-1D-Single’s outputs (Singh *et al.*, 2021b) outperform the single-sequence-based SSCpred and the MSA-based predictors for proteins with a low effective number of homologous proteins (Neff). We also showed that an ensemble of models trained through different training approaches and different feature combinations adds to this improvement.

## 2 Materials and methods

### 2.1 Datasets

To curate a dataset, we utilized the benchmark dataset prepared by ProteinNet (AlQuraishi, 2019). It consists of 50914 proteins submitted to PDB before 2016 with high resolution (< 2.5Å) crystal structures and clustered at sequence identity cut-off at 95% according to MMseqs2 tool (Steinegger and Söding, 2017). ProteinNet provides a number of datasets at different sequence identity cut-offs, but we chose the dataset with the sequence identity cut-off of 95% for training to obtain as much data as possible to harness the full capabilities of recent deep learning algorithms.

To efficiently validate models during training and minimize possible over-fitting, we separated 100 proteins from the ProteinNet set and compared their Hidden Markov Models generated by HHblits with the Hidden Markov Models of other proteins in the training dataset and validation set using HHblits. Any proteins, which had hits with these 100 validation proteins at an e-value cut-off of less than 0.1, were removed. This left us with the final 39120 proteins for the training set. After removing any proteins with a length more than 500 from both the training and validation sets, the final training and validation sets have 34691 and 88 proteins, respectively.

For independent testing and comparison, we downloaded all protein structures released between May 2018 and April 2020. As it can be insufficient to remove homologous sequences, we removed any potential homologs in the training set to the test data by comparing the Hidden Markov Models of all post-2018 proteins to the Hidden Markov Models of all pre-2018 proteins using the HHblits tool at an e-value cut-off of less than 0.1 (Steinegger *et al.*, 2019a). This led to 669 proteins as a stringent test set named SPOT-2018.

To test how predictors perform on de-novo proteins and proteins without homologs, we separated 46 proteins from SPOT-2018 which have Neff=1 forming a test set called Neff1-2018. This provides a reliable, stringent and completely independent benchmark to compare the performance of different predictors on sequentially isolated proteins. Neff is calculated with respect to the reference Uniclust30 dataset (Published Feb 2020).

Apart from SPOT-2018 and Neff1-2018, we employed an additional independent test set CASP14-FM. This test set includes 15 free modelling targets released at CASP14 (Liu *et al.*, 2021). Free modeling targets are those proteins without known structural templates in the protein databank at the time of release. Supplementary Table S1 provides a brief description of the test sets utilized in this study.

### 2.2 Input features

To train an ensemble of neural networks proposed in this method, we used multiple combinations of several features, including one-hot encoding of amino acids, the output of SPOT-1D-Single and attention maps from ESM-1b (Rao *et al.*, 2020). One-dimensional features of one-hot encoding and the output of SPOT-1D-Single were converted into two-dimensional features using an outer concatenation. From SPOT-1D-Single, we used the probabilities of three-state-secondary-structure (SS3) and eight-state-secondary structure (SS8), Solvent Accessible Surface Area (ASA), Half-Sphere-Exposure (HSE), and protein backbone torsion angles *ψ*, *ϕ*, *θ* and *τ*. Attention maps from ESM-1b were gathered by using all twenty attention heads from the last layer of the transformer as well as twenty attention heads from every layer of the ESM-1b model. For both cases, we symmetrized and applied average product corrections (APC) to the extracted attention maps as done by (Rao *et al.*, 2020).

### 2.3 Performance evaluation

The aim of this research is to predict which amino acid pairs in a protein are in contact. Following the standard CASP definition (Ezkurdia *et al.*, 2009), protein residues are considered to be in contact when there is an inter-residue distance of ≤ 8.0 Å between two *C_β_* atoms. A contact between two residues is classified into three types: long (at least 24 residues apart), medium (between 12 to 23 residues apart) and short (between 7-11 residues apart) ranges. For these three types of contacts, we calculated top L/10, L/5, L/2, and L/1 highest-ranked predictions in terms of precision. For further assessment in this work, we also calculated the overall F1-score, Matthews Correlation Coefficient (MCC) (Chicco and Jurman, 2020), Sensitivity, Area Under Curve of Precision-Recall Curve (AUC), and Area Under Curve of Receiver Operating Characteristic (ROC) of short-, medium-, and long-range contacts, together. We also obtained the F1-score, MCC, sensitivity for our model and all other predictors at the maximum F1-score cut-off for the data set.

### 2.4 Neural Networks

Our deep neural network architecture was inspired by the success of the ResNet architecture in protein contact-map and RNA secondary structure prediction (Wang *et al.*, 2017;Singh *et al.*, 2021a). In this paper, we use a 12 block ResNet, which is the maximum depth that we could train on the available GPU. Instead of using vanilla ResNet models, we employed a recently published version of ResNet (Duta *et al.*, 2020). This improved version of ResNet was shown to perform better than vanilla and pre-act ResNet for both image and video-based tasks. Here, we applied this architecture for the inter-residue contact prediction problem.

As shown in Fig. 1, we employed convolutional layers with a channel size of 64 and kernel size of 3. We trained six models with the same architectural specifications but different input feature combinations as described in Table 1. The first three models in Table 1 were trained to predict the inter-residue contacts as a binary classification, while for the last three models, we predicted inter-residue distances as distance bins, and then we added the probabilities within the bins for the distances between 0-8 Å.

**Fig. 1:**
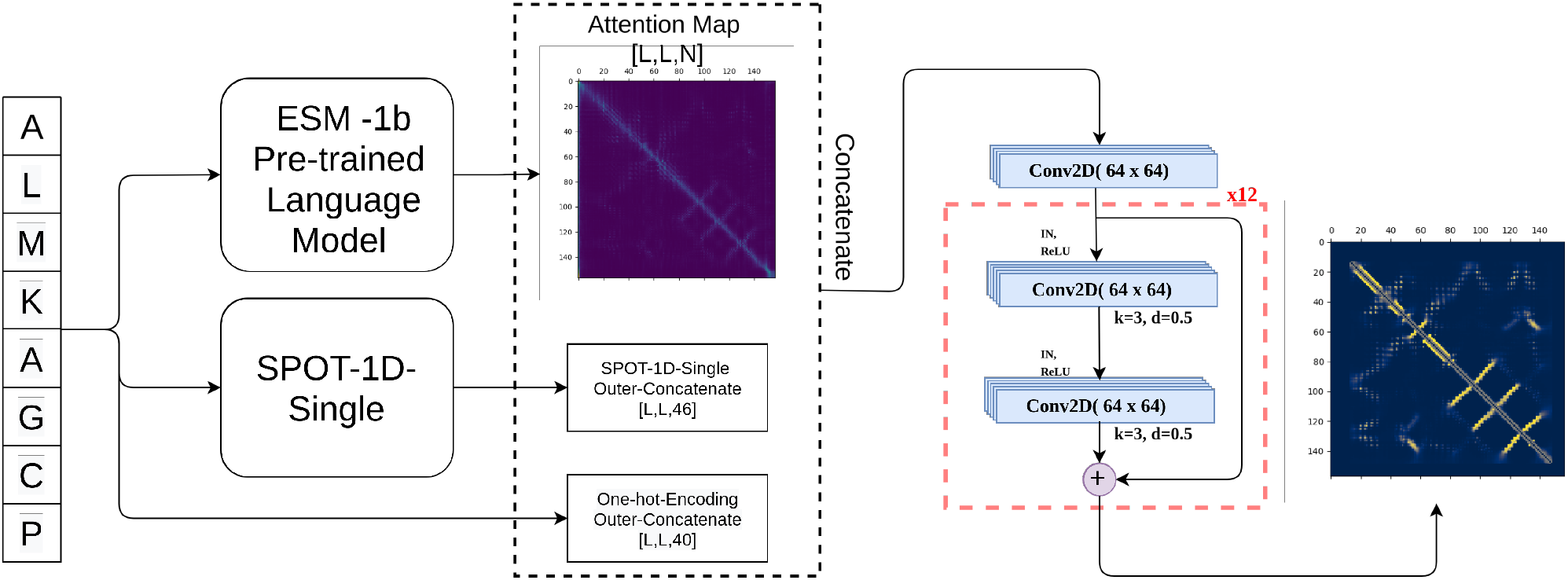
Overview of the model pipeline.

**Table 1.**
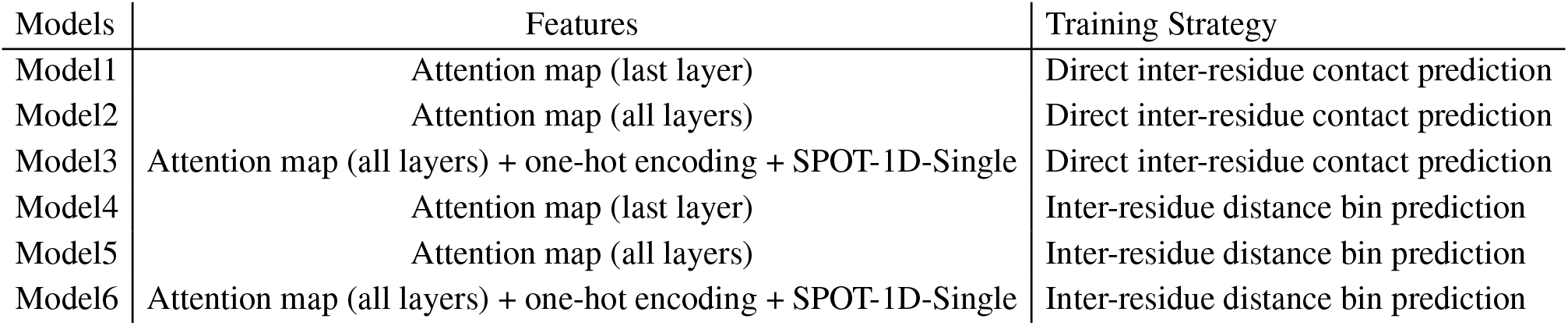
A description of feature combinations for the ensemble of trained models.

| Models | Features | Training Strategy |
| --- | --- | --- |
| Model1 | Attention map (last layer) | Direct inter-residue contact prediction |
| Model2 | Attention map (all layers) | Direct inter-residue contact prediction |
| Model3 | Attention map (all layers) + one-hot encoding + SPOT-1D-Single | Direct inter-residue contact prediction |
| Model4 | Attention map (last layer) | Inter-residue distance bin prediction |
| Model5 | Attention map (all layers) | Inter-residue distance bin prediction |
| Model6 | Attention map (all layers) + one-hot encoding + SPOT-1D-Single | Inter-residue distance bin prediction |

The direct contact-map prediction models were trained using Binary Cross-Entropy loss, while the distogram-based prediction models were trained using Cross-Entropy loss. Apart from this major difference, other model hyperparameters and specifications are the same. This includes using the Adam optimizer with a learning rate of 0.001 and a batch size of 1. To avoid overfitting, all models were trained with early stopping of 3.

### 2.5 Method comparison

We compared SPOT-Contact-Single with language model’s supervised regression contact-map predictor ESM-1b, single-sequence-based SSCpred, profile-based-predictors TrRosetta and SPOT-Contact. The above-stated methods TrRosetta, SPOT-Contact and ESM-1b have stand-alone programs available online from https://github.com/gjoni/trRosetta, https://sparks-lab.org/server/spot-contact/, and https://github.com/facebookresearch/esm, respectively. Input to all profile-based methods including TrRosetta was obtained from SPOT-Contact MSA generation pipeline for benchmarking purposes. For SSCpred, we utilized the web-server available online from http://csbio.njust.edu.cn/bioinf/sscpred/ due to lack of its stand-alone version.

## 3 Results

### 3.1 Feature importance

To understand the effect of different features, we trained a ResNet12 architecture on different input features and compared their performance on the validation set. Table 2 shows that the model trained on the one-hot encoding of the fasta sequence only predicts the contact-map with an F1-score of 0.15 and an MCC of 0.14. Adding the output of SPOT-1D-Single (a single-sequence-based predictor) to one-hot encoding improved the F1-score by 22.90%. By comparison, using the attention map output from the unsupervised learning method, ESM-1b significantly boosted the performance. The attention maps extracted from the last layer of the ESM-1b lead to 0.51, 0.50, 0.56, and 0.95 for F1, MCC, precision and AUC of ROC, respectively. The result is 242% and 178% improvement in the F1-score over models trained on one-hot encoding and SPOT-1D-Single + one-hot encoding, respectively. Using the attention maps extracted from all layers of ESM-1b, Table 2 shows similar results for F1-score and MCC as the attention map from the last layer, but with continued improvement in the precision of all short-, medium-, and long-range contacts (L/10, L/5, L/2) (Supplementary Table S2). As expected, concatenating all features together (one-hot encoding + SPOT-1D-Single + ESM-1B attention maps (all layers)) further showed an increase of 2.92% in F1-score over using the attention maps (all layers) only with noticeable improvement in model precision for medium- and long-range contacts, in particular (Supplementary Table S2). Based on these results, we further trained different model architectures using a combination of one-hot encoding, SPOT-1D-Single and ESM-1b attention maps (all layers).

**Table 2.** Comparison of ResNet12’s model performance trained on different feature combinations on the validation set. We compared the F1-score, MCC, Sensitivity, Precision, AUC, and ROC for predicting all contacts (short-, medium-, and long-range).

|  | Feature | F1 | MCC | Sensitivity | Precision | AUC | ROC |
| --- | --- | --- | --- | --- | --- | --- | --- |
| 1 | One-hot encoding | 0.1489 | 0.1365 | 0.1815 | 0.1261 | 0.0774 | 0.7421 |
| 2 | One-hot encoding + SPOT-1D-Single | 0.1830 | 0.1670 | 0.1863 | 0.1799 | 0.1169 | 0.7427 |
| 3 | ESM-1b attention map (last layer only) | 0.5095 | 0.5031 | 0.4663 | 0.5615 | 0.5167 | 0.9471 |
| 4 | ESM-1b attention map (all layers) | 0.5098 | 0.5011 | 0.4897 | 0.5315 | 0.5199 | 0.9481 |
| 5 | All features | 0.5244 | 0.5189 | 0.4758 | 0.5840 | 0.5382 | 0.9492 |

### 3.2 Direct vs distance contact-map prediction

To predict protein contact maps, we examined two different training strategies: direct contact-map prediction and distogram-based contact-map prediction by training a ResNet12 on one-hot encoding, SPOT-1D-Single’s output and ESM-1b’s attention maps concatenated together. Table 3 and Supplementary Table S3 shows that direct contact-map prediction performs slightly better, but the difference between the two training strategies is small. Thus, both strategies were employed in different models for our final ensemble.

**Table 3.** Performance comparison of two training strategies: direct contact- and distogram contact-map prediction on the validation set.

| Model | F1 | MCC | Sensitivity | Precision | AUC | ROC |
| --- | --- | --- | --- | --- | --- | --- |
| Direct Contact Prediction | 0.5244 | 0.5189 | 0.4758 | 0.5840 | 0.5382 | 0.9492 |
| Distogram Contact Prediction | 0.5201 | 0.5128 | 0.4859 | 0.5596 | 0.5264 | 0.9458 |

### 3.3 Ensemble learning performance

Based on the findings of the previous two sections, we trained six different models with three best feature combinations using both distogram and direct contact prediction. We then ensembled the results of all six models to gain improvement over individual models by taking the mean of individual models. To understand the improvement gained, Table 4 presents the results of the selected six individual models and the ensemble of the six models on the validation set. The performance of the ensemble (SPOT-Contact-Single) is higher than all individual models, with 4% improvement over the second-best individual model for the F1-score. This performance gain is consistent across all matrices.

**Table 4.** Individual model performance as compared to the ensemble performance on the validation set for contact-map prediction.

| Model | F1 | MCC | Sensitivity | Precision | AUC | ROC |
| --- | --- | --- | --- | --- | --- | --- |
| Model1 | 0.5095 | 0.5031 | 0.4663 | 0.5615 | 0.5167 | 0.9471 |
| Model2 | 0.5098 | 0.5011 | 0.4897 | 0.5315 | 0.5199 | 0.9481 |
| Model3 | 0.5244 | 0.5189 | 0.4758 | 0.5840 | 0.5382 | 0.9492 |
| Model4 | 0.5201 | 0.5128 | 0.4859 | 0.5596 | 0.5264 | 0.9458 |
| Model5 | 0.5188 | 0.5123 | 0.4770 | 0.5686 | 0.5246 | 0.9396 |
| Model6 | 0.5117 | 0.5045 | 0.4750 | 0.5545 | 0.5108 | 0.9423 |
| SPOT-Contact-Single | 0.5444 | 0.5389 | 0.4977 | 0.6008 | 0.5664 | 0.9586 |

### 3.4 Method comparison

Because our method does not employ evolutionary information, it is fair to compare all methods (evolution-based and single-sequence-based) on the proteins without homologous sequences. Table 5 compares the performance of SPOT-Contact-Single (this work) with ESM-1b (LM), SPOT-Contact (evolution-based) and TrRosetta (evolution-based) for those proteins with Neff=1 in the SPOT-2018 set (Neff1-2018). The evolution-based techniques (SPOT-Contact and TrRosetta) achieve similar performance as ESM-1b with F1 scores at about 0.22. By comparison, the F1-score given by SPOT-Contact-Single is 45% higher at 0.32. Similar trends are observed across other performance measures, including the precision for top predictions at short-, medium-, and long-range contacts, as shown in Fig. 2.

**Table 5.**
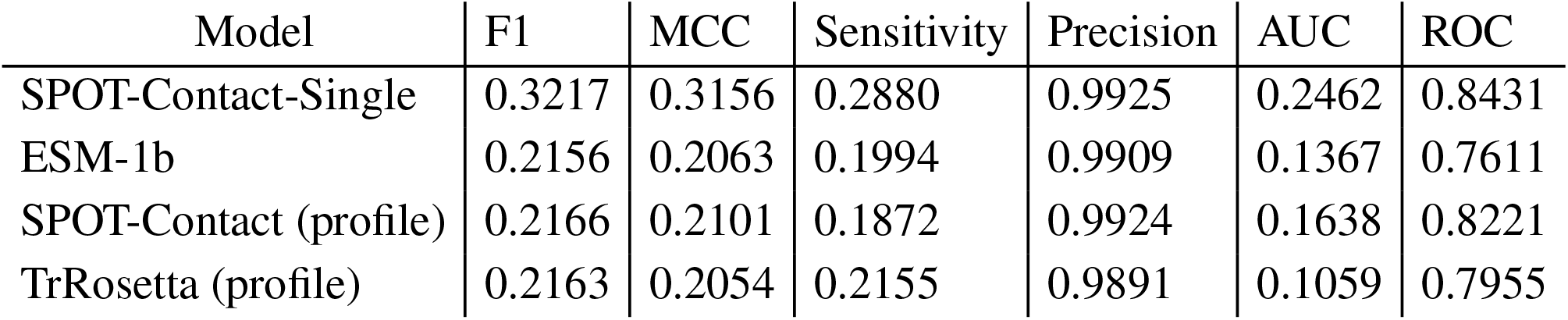
Comparison of SPOT-Contact-Single, SPOT-Contact, TrRosetta, and esm-1b on the Neff=1 set (Neff1-2018). To measure the performance of the predictors, we compared the F1-score, MCC, Precision, Sensitivity, AUC, and ROC of the overall prediction for all short-, medium-, and long-range predictions collectively, for the highest threshold of each predictor for this test set.

**Fig. 2:**
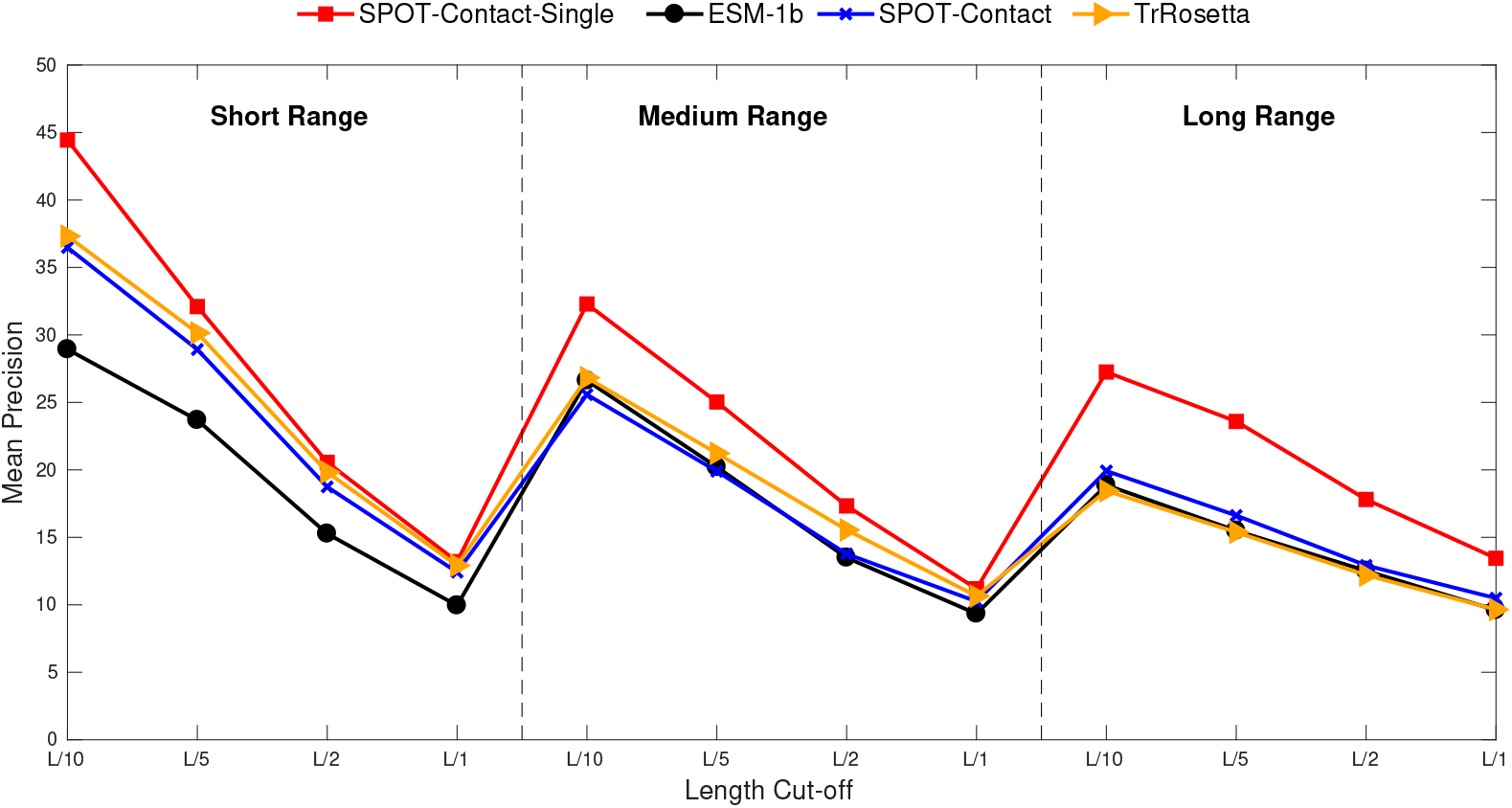
Precision-based comparison of SPOT-Contact-Single, SPOT-Contact, TrRosetta, and ESM-1b on Neff1-2018 for short-, medium-, and long-range contacts.

To illustrate the effect of homologous sequences, we plotted the F1-score of different predictors as a function of the Neff values in Fig. 3. The performance of the profile-based predictors improves over SPOT-1D-Single as Neff increases. In other words, SPOT-1D-Single is not yet as competitive as evolution-based methods. This is because multiple sequence alignment of homologous sequence can provide co-mutation information more effectively than unsupervised learning.

**Fig. 3:**
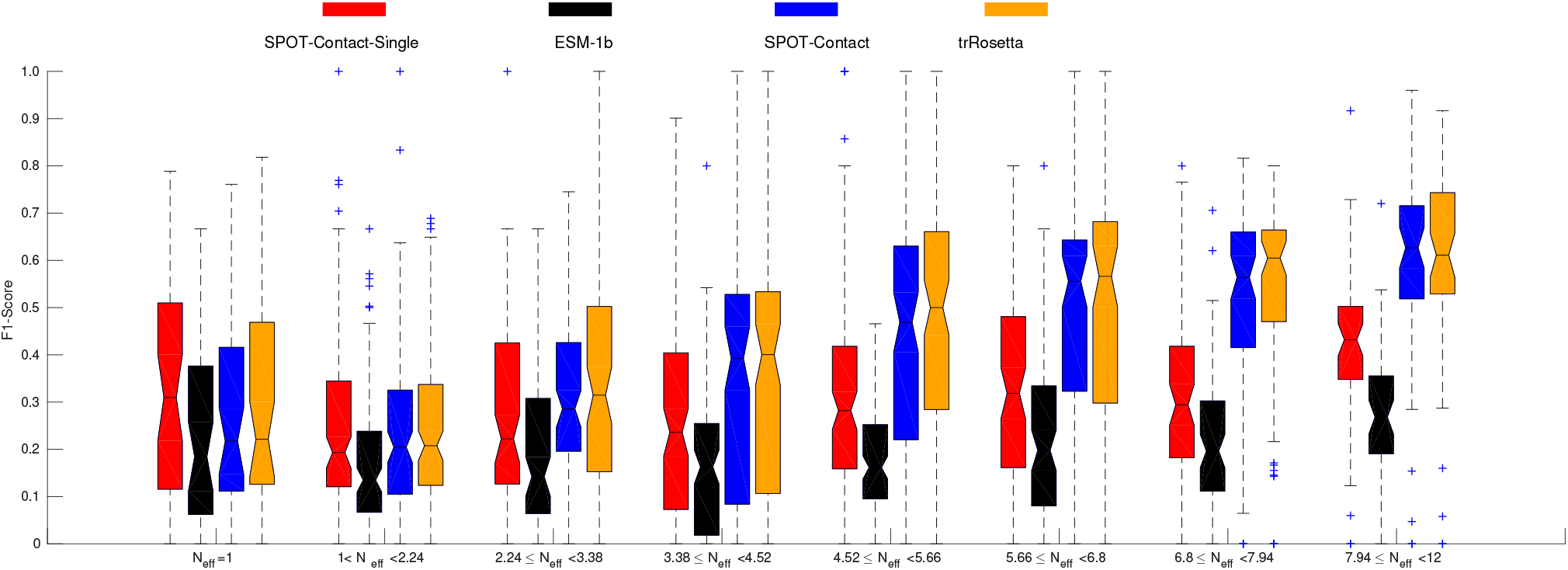
F1-score as a function of the number of effective homologous sequences (Neff) by SPOT-Contact-Single compared with other methods on SPOT-2018 for contact-map prediction.

The native and predicted contact-maps from SPOT-Contact-Single, SPOT-Contact, TrRosetta and ESM-1b on an example protein (5YKZ_A) from Neff1-2018 are presented in Fig. 4, which shows SPOT-Contact-Single provided a more accurate prediction of the contact-map for this low Neff protein, with the F1-scores of 0.215, 0.235, 0.252, and 0.388 for SPOT-Contact, TrRosetta, ESM-1b, and SPOT-Contact-Single, respectively.

**Fig. 4:**
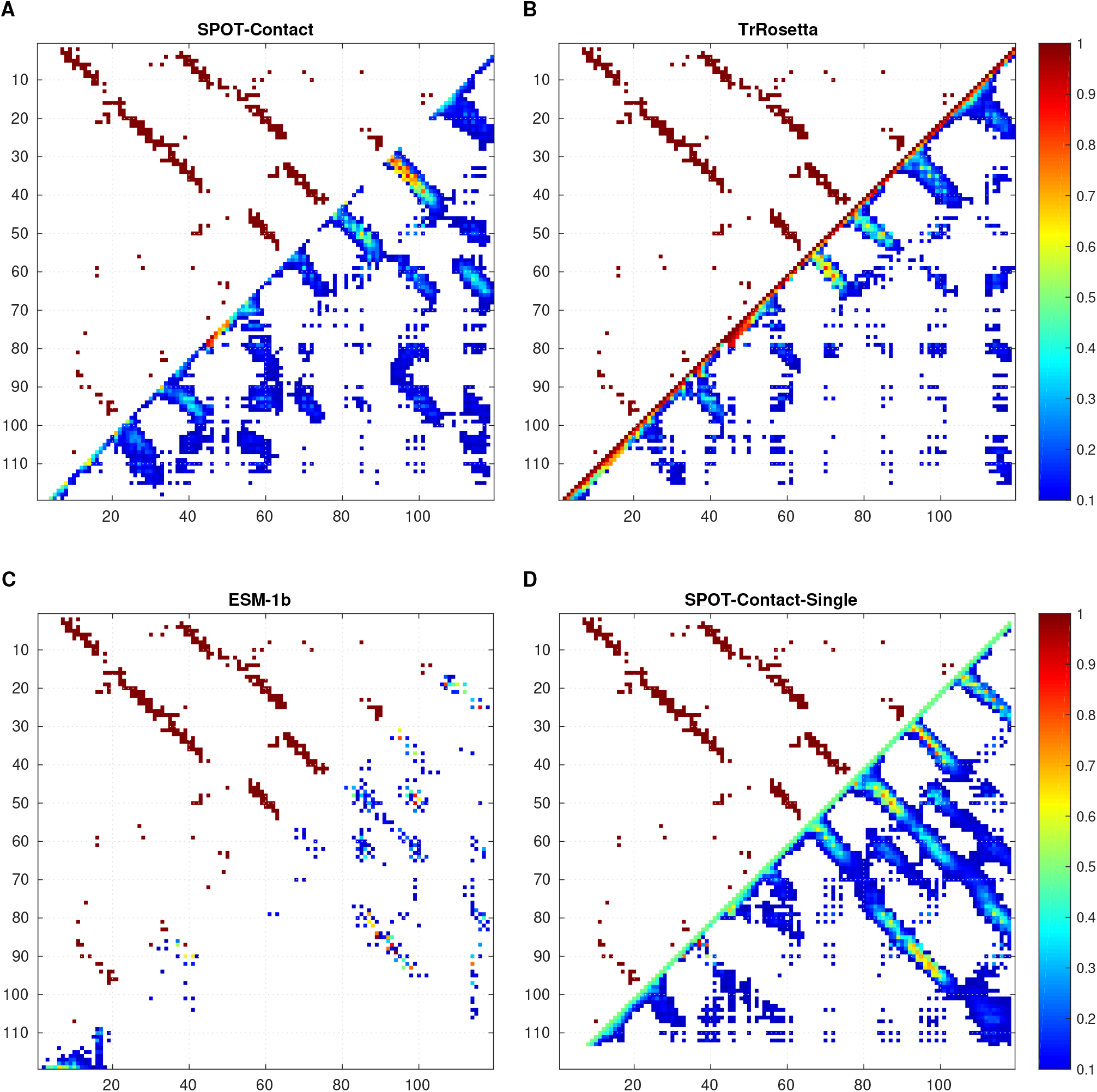
Comparison of the predictions for 5YKZ_A protein by four methods as labeled. The upper triangle and lower triangle represent the native and the predicted contact-map, respectively.

### 3.5 Comparison with SSCpred

SSCpred is also a single-sequence-based contact-map predictor that employed the proteins released till 2019 April for training. To make a fair comparison, we compared SSCpred to other predictors on the CASP14-FM dataset. Table 6 shows that SPOT-Contact-Single performs much better than ESM-1b and SSCpred according to F1-score (15% improvement over SSCpred) and other measures. The most improvement over SSCpred is in long-range contacts precision (~100% improvement, Supplementary Table S4).

**Table 6.** Comparison of SPOT-Contact-Single, SSCpred, and ESM-1b on CASP14-FM. To measure the performance of the predictors, we compared the F1-score, MCC, Precision, Sensitivity, AUC, and ROC of the overall prediction for all short-, medium-, and long-range predictions collectively, for the highest threshold of each predictor for this test set.

| Model | F1 | MCC | Sensitivity | Precision | AUC | ROC |
| --- | --- | --- | --- | --- | --- | --- |
| SPOT-Contact-Single | 0.2061 | 0.1925 | 0.2069 | 0.2053 | 0.1322 | 0.7953 |
| SSCpred | 0.1797 | 0.1651 | 0.1943 | 0.1671 | 0.1088 | 0.7772 |
| ESM-1b | 0.1513 | 0.1372 | 0.1482 | 0.1546 | 0.0765 | 0.6776 |

## 4 Discussion

In this paper, we have developed a new protein contact-map predictor which employs the pretrained features from a transformer language model as input to predict contact maps without using homologous sequences. We employed an ensemble of ResNet-based architectures trained on multiple combinations of several features and a large training set of almost 35000 proteins with validation and test sets that are non-redundant to all training proteins according to HHsearch. The accuracy of SPOT-Contact-Single is higher than the evolutionary-profile-based SPOT-1D and TrRosetta when the number of effective homologous sequence is low. This highlights that SPOT-Contact-Single can be used as a reasonably accurate screening tool for protein contact map prediction.

Using ESM-1b attention map in SPOT-Contact-Single makes it not possible to directly predict contact maps for proteins with more than 1024 amino acids. This should not prevent the use of SPOT-Contact-Single for large proteins because proteins are usually made of domains with less than 1000 residues.

A point of interest could be to profile our method (SPOT-Contact-Single) against a profile-based method (TrRosetta) in terms of computational time. As shown in Supplementary Table S5, while running inference on CPU for CASP14-FM dataset of 15 proteins, SPOT-Contact-Single makes the prediction in 116 seconds which is 22 times faster than TrRosetta. Also, on GPU, TrRosetta took 1926 seconds which 42 times slower than SPOT-Contact-Single. As expected, the sequence profile generation takes significantly longer than the proposed method making the latter more suitable for genomic scale prediction.

Finally, SPOT-Contact-Single predicts the protein contact-map without using evolutionary features. The further improvement in protein contact-map prediction without evolutionary information may come from using more advanced architectural models such as Transformer (Vaswani *et al.*, 2017) or Performer (Choromanski *et al.*, 2020) for downstream supervised training.

## Acknowledgements

We gratefully acknowledge the use of the High Performance Computing Cluster Gowonda to complete this research, and the aid of the research cloud resources provided by the Queensland Cyber Infrastructure Foundation (QCIF). We also gratefully acknowledge the supercomputing facility at the Shenzhen Bay Laboratory and the support of NVIDIA Corporation with the donation of the Titan V GPU used for this research.

