## Supplementary material for "SPOT-Contact-Single: Improving Single-Sequence-Based Prediction of Protein Contact Map using a Transformer Language Model"

**Supplementary Table S1:** A description of different test sets used in this research.

| Dataset Name | Protein Count | Validation tool / Body | E-value Cutoff | Description | Average Neff |
|---|---|---|---|---|---|
| SPOT-2018 | 669 | HHsearch | 0.01 | Post 2018 proteins independent to all pre-2018 proteins | 4.47 |
| Neff1-2018 | 46 | HHsearch | 0.01 | SPOT-2018 subset of proteins with no homologs | 1.0 |
| CASP14-FM | 15 | CASP targets | - | Free modelling targets released during CASP-14 | 2.43 |

**Supplementary Table S2:** Comparison of model precisions by using ResNet12 trained on different feature combinations for short-, medium-, and long-range contacts on the validation set.

| | Model | Short Range Contacts | | | | Medium Range Contacts | | | | Long Range Contacts | | | |
|---|---|---|---|---|---|---|---|---|---|---|---|---|---|
| | | L/10 | L/5 | L/2 | L/1 | L/10 | L/5 | L/2 | L/1 | L/10 | L/5 | L/2 | L/1 |
| 1 | One-hot encoding | 28.82 | 23.94 | 17.33 | 12.88 | 19.73 | 16.78 | 12.84 | 10.14 | 7.66 | 6.63 | 5.72 | 4.91 |
| 2 | One-hot encoding + SPOT-1D-Single | 35.29 | 30.53 | 27.73 | 24.91 | 20.46 | 17.25 | 12.23 | 10.98 | 10.69 | 9.93 | 8.34 | 5.95 |
| 3 | ESM-1b attention map (last layer only) | 76.99 | 63.51 | 38.93 | 23.23 | 77.20 | 66.27 | 46.43 | 29.17 | 83.45 | 78.49 | 65.95 | 51.81 |
| 4 | ESM-1b attention map (all layers) | 78.53 | 64.23 | 39.10 | 23.18 | 77.69 | 67.23 | 45.57 | 28.80 | 83.59 | 78.66 | 66.23 | 52.22 |
| 5 | All features | 78.39 | 63.86 | 39.04 | 23.35 | 79.51 | 68.77 | 47.00 | 29.66 | 85.76 | 81.31 | 68.85 | 54.16 |

**Supplementary Table S3:** Precision comparison of two training strategies: direct contact prediction, and distogram contact prediction for short-, medium-, and long-range contacts on the validation set.

| Model | Short Range Contacts | | | | Medium Range Contacts | | | | Long Range Contacts | | | |
|---|---|---|---|---|---|---|---|---|---|---|---|---|
| | L/10 | L/5 | L/2 | L/1 | L/10 | L/5 | L/2 | L/1 | L/10 | L/5 | L/2 | L/1 |
| Direct Contact Prediction | 78.39 | 63.86 | 39.04 | 23.35 | 79.51 | 68.77 | 47.00 | 29.66 | 85.76 | 81.31 | 68.85 | 54.16 |
| Distogram Contact Prediction | 76.22 | 63.38 | 39.09 | 23.63 | 76.15 | 67.09 | 46.82 | 29.73 | 84.15 | 78.49 | 67.19 | 53.65 |

**Supplementary Table S4:** Precision-based comparison of SPOT-Contact-Single, SSCpred, SPOT-Contact, TrRosetta, and ESM-1b on the CASP14-FM set for short, medium and long range contacts.

| Model | Short Range Contacts | | | | Medium Range Contacts | | | | Long Range Contacts | | | |
|---|---|---|---|---|---|---|---|---|---|---|---|---|
| | L/10 | L/5 | L/2 | L/1 | L/10 | L/5 | L/2 | L/1 | L/10 | L/5 | L/2 | L/1 |
| SPOT-Contact-Single | 47.30 | 39.64 | 26.14 | 18.67 | 29.73 | 24.72 | 17.96 | 13.93 | 18.92 | 19.38 | 15.40 | 11.56 |
| SSCpred | 37.84 | 33.18 | 24.82 | 18.72 | 26.13 | 24.50 | 17.17 | 12.61 | 9.91 | 9.13 | 7.66 | 7.69 |
| ESM-1b | 35.14 | 28.29 | 18.13 | 13.22 | 22.97 | 19.82 | 15.58 | 10.85 | 17.12 | 12.47 | 9.42 | 7.38 |

**Supplementary Table S5:** Inference time comparison of SPOT-Contact-Single and TrRosetta for prediction on 15 proteins of CASP14-FM.

| Computational Specifications | TrRosetta | SPOT-Contact-Single |
|---|---|---|
| 48 CPU threads on Intel(R) Xeon(R) CPU E5-2670 v3 @ 2.30GHz | 2576 Seconds | 131 Seconds |
| TITAN X (Pascal) | 1926 Seconds | 46 Seconds |
